# Individualized surface parcellation enhances characterization of resting-state brain dynamics and their alterations in schizophrenia

**DOI:** 10.64898/2026.07.24.740570

**Authors:** Harrison Watters, Petra Fürstová, Jaroslav Tintěra, Filip Spaniel, Jaroslav Hlinka

## Abstract

Resting-state functional MRI (rs-fMRI) studies in schizophrenia commonly rely on normalization to volumetric templates and fixed atlas parcellations derived from neurotypical populations. While these approaches enable group-level comparisons, they may obscure individual variation in cortical organization and intrinsic brain dynamics. In this study, we compared four preprocessing and parcellation strategies across two independent schizophrenia cohorts (MRI site 1, n=159; MRI site 2, n=255) to evaluate how analytic choices affect static functional connectivity and dynamic quasi-periodic pattern (QPP) measures, including default mode-dorsal attention network opposition, QPP component rank, explained variance, event rate, and associations with PANSS symptom severity.

Across datasets, individualized surface-based parcellation (IndiPar) consistently detected more pronounced QPP dynamics, stronger default mode / dorsal attention network opposition, and greater explained variance of QPPs relative to atlas-based pipelines. IndiPar also produced larger and more reproducible patient-control differences in functional connectivity and QPP event-rate measures, suggesting improved sensitivity through preservation of subject-specific organization. IndiPar additionally detected a significantly increased QPP event rate and more symptom associations in patients in the larger dataset. However, associations between fMRI measures and symptom severity showed limited stability across cohorts.

These findings extend previous reports of altered resting-state activity in schizophrenia, and demonstrate that preprocessing and parcellation choices substantially influence both static and dynamic rs-fMRI results. Individualized surface-based parcellation appears to better preserve subject-specific variability and improves detection of intrinsic brain dynamics. At the same time, the limited cross-dataset replication of symptom associations highlights the challenges of deriving stable brain-symptom relationships from heterogeneous psychiatric cohorts.

## 1. Introduction

Brain abnormalities have been widely reported in the Schizophrenia neuroimaging literature (Mwansisya, 2017; Dabiri, 2022; Voineskos, 2024). However, most analyses focus on group-level averages, and these findings have rarely translated into clinically useful biomarkers (Woo, 2018; Kraguljac, 2021). As a result, a large gap remains between the fields of psychiatry and neuroimaging (Cuthbert, 2026).

Part of the problem is that group-level analyses presume there exists a set of unifying features or tendencies explaining each of our categorical definitions of psychiatric disorders - a bottleneck beneath the noise perhaps - despite significant individual variability and heterogeneity of symptoms (Wolfers, 2018; Habtewold, 2020). But there is growing recognition in the field that our diagnostic categories of psychiatric disorders stretch far beyond empirical evidence (Kotov, 2021; Voineskos, 2022). Findings from neuroimaging increasingly suggest that individual differences in functional brain organization are stable, reproducible, and behaviorally relevant rather than simply measurement noise (Miller, 2012; Mueller, 2013). Indeed, even in neurotypical subjects, inter-individual variation in intrinsic functional activity is consistent across task/rest conditions and scan sessions, and is, albeit far from perfectly, predictive of cognitive measures such as fluid intelligence (Finn, 2015).

Variation is especially important in psychiatric disorders such as schizophrenia spectrum disorder, where patients show substantial heterogeneity in symptom profiles, illness trajectories, genetic risk, and functional and structural brain alterations, alongside considerable genetic overlap with other psychiatric disorders (Brugger, 2017; Smeland, 2020; Fang, 2022). Heterogeneity in psychiatric disorders, and incompatibility of empirical neuroimaging findings with legacy diagnostic categories, is such that the whole approach of categorical psychiatry has come into question (Insel, 2010; Kotov, 2017; Hengartner, 2017; Kotov, 2021), with dimensional psychiatry frameworks such as HiTOP (Kotov, 2021) and RDoC (Insel, 2010) proposed as possible alternatives.

Standard pre-processing practices in functional MRI have likely contributed to shortcomings in translational neuroimaging. The goal of two of the most commonplace fMRI preprocessing steps (standardization to a volumetric template space such as MNI-152, and parcellation of voxels into an atlas space) is to permit comparison between brains. However, as mentioned, no two brains are alike, and while such standardization strategies permit greater comparison, they do not ensure such comparisons are apples-to-apples or that they reflect biological reality (Shifferman, 2015). Imposing normalization and identical parcellation schemes across subjects may obscure meaningful variation in functional and anatomical connectivity. The more neurodivergent the population is, the more this ought to concern us.

To address the limitations of standardized parcellation approaches in fMRI, previously, Wang and colleagues developed an individual-specific parcellation approach for use in functional MRI time series, which allows for the generation of individualized parcellations and homologous functional region identification (Wang, 2015). This approach is conducted on surface-based time series which retain a greater degree of subjects’ unique neocortical brain topography. Using this method, they found improved estimation of SZ symptom scores (PANSS scores) versus functional connectivity (Wang, 2019) compared to standard atlas parcellation approaches.

Quasi-Periodic Patterns (QPPs) are recurring infraslow (∼0.05-0.1 Hz) patterns of large-scale brain activity characterized by alternating engagement of task-positive and task-negative networks (Majeed, 2011). These intrinsic, weather-like spatiotemporal dynamics can be reliably detected in resting and task scans (Watters, 2025) using sliding-window approaches (Xu, 2025) and complex principal components analysis (cPCA) (Bolt, 2022). Although QPPs involve distributed large-scale network activity, their most prominent and consistently reported feature is alternating opposition between the default mode network (DMN) and dorsal attention network (DAN) (Bolt, 2022; Abbas, 2019b). There is evidence that the balance of activity between these networks is important for attentional processing and cognitive state regulation (Abbas, 2019b; Seeburger, 2024). Because schizophrenia has repeatedly been associated with altered DMN function and internally directed cognition, DMN-related QPP dynamics may be particularly relevant in this population (Raichle, 2015; Menon, 2023; Halari, 2009; Thomas, 2014). Indeed, prior work suggests that QPP-related fluctuations may contribute more strongly to intrinsic functional connectivity in individuals with schizophrenia compared with healthy controls (Briend, 2020).

In addition to computing functional connectivity and QPP events using IndiPar, we compare the IndiPar approach results directly to three more traditional pre-processing approaches: 1) Surface-based time series in standardized Schaefer 200 atlas space, 2) Volumetric normalized to MNI-152 in Schaefer 200 atlas space, and 3) Volumetric normalized to MNI-152 in Brainnetome 246 atlas space.

Results from single-dataset SZ studies often fail to generalize between sites and cohorts, even in studies employing machine-learning approaches (Orban, 2018; Eitel, 2021; Li, 2025), with some inter-site classification rates falling to essentially chance levels (Cai, 2020). To prevent over-inference on results from a single-dataset, we analyzed the 4 fMRI-pipelines in two independent cohorts of SZ patients from separate sites to evaluate generalizability of results between datasets. All analyses were conducted at the subject level before assigning patient vs control groups to avoid group-level logic and ecological fallacy (Subramanian, 2009).

The primary hypotheses were the following:

1. IndiPar should more robustly detect canonical quasi-periodic patterns (QPPs), reflected by stronger DMN-DAN opposition, greater explained variance, and higher prevalence of rank-1 QPP components, because surface-based individualized parcellation better preserves subject-specific cortical topography.
2. Individualized parcellation should produce greater similarity between cohorts (IKEM vs NUDZ) and improve sensitivity to patient-control differences, as it may reduce variability introduced by scanner differences and volumetric normalization to template spaces such as MNI-152.
3. Consistent with Wang et al., 2019, the IndiPar approach should preserve stronger associations between functional imaging measures (static and dynamic in this case) and patient symptom scores (PANSS) relative to atlas-based approaches.

The present study systematically examines how preprocessing and parcellation choices influence both static and dynamic resting-state fMRI findings in schizophrenia. Using two independent cohorts, we compare an individualized surface-based parcellation framework (IndiPar) with three commonly used atlas-based pipelines and extend this comparison to infraslow dynamic activity captured by quasi-periodic patterns (QPPs). By evaluating patient-control differences, cross-dataset reproducibility, and associations with symptom severity across pipelines, this study tests whether individualized surface-based approaches better preserve subject-specific neural variability and improve the detectability of disease-specific alterations in intrinsic brain dynamics.

## 2. Methods

### 2.1 Participants and Datasets

Resting-state functional MRI (rs-fMRI) data were analyzed from two independent cohorts of first-episode schizophrenia patients and healthy controls collected at separate MRI sites in the Czech Republic: the Institute for Clinical and Experimental Medicine (IKEM) and the National Institute of Mental Health (NUDZ). De-identified scans were processed and analyzed at the Czech Academy of Sciences, Institute of Computer Science.

The final sample included 159 participants from IKEM (92 schizophrenia patients, 67 controls) and 255 participants from NUDZ (153 schizophrenia patients, 102 controls, see Figure 1 for age and sex demographics). Patients were diagnosed according to ICD-10 clinical criteria during the early phase of treatment with second-generation antipsychotic medication. Healthy controls were included only if they had no lifetime history of major psychiatric disorders or family history of psychotic disorders. Exclusion criteria for all participants included history of neurological illness, significant head trauma, substance dependence, or contraindications to MRI. All participants provided written informed consent in accordance with institutional review board approvals at both sites.

**Figure 1:**
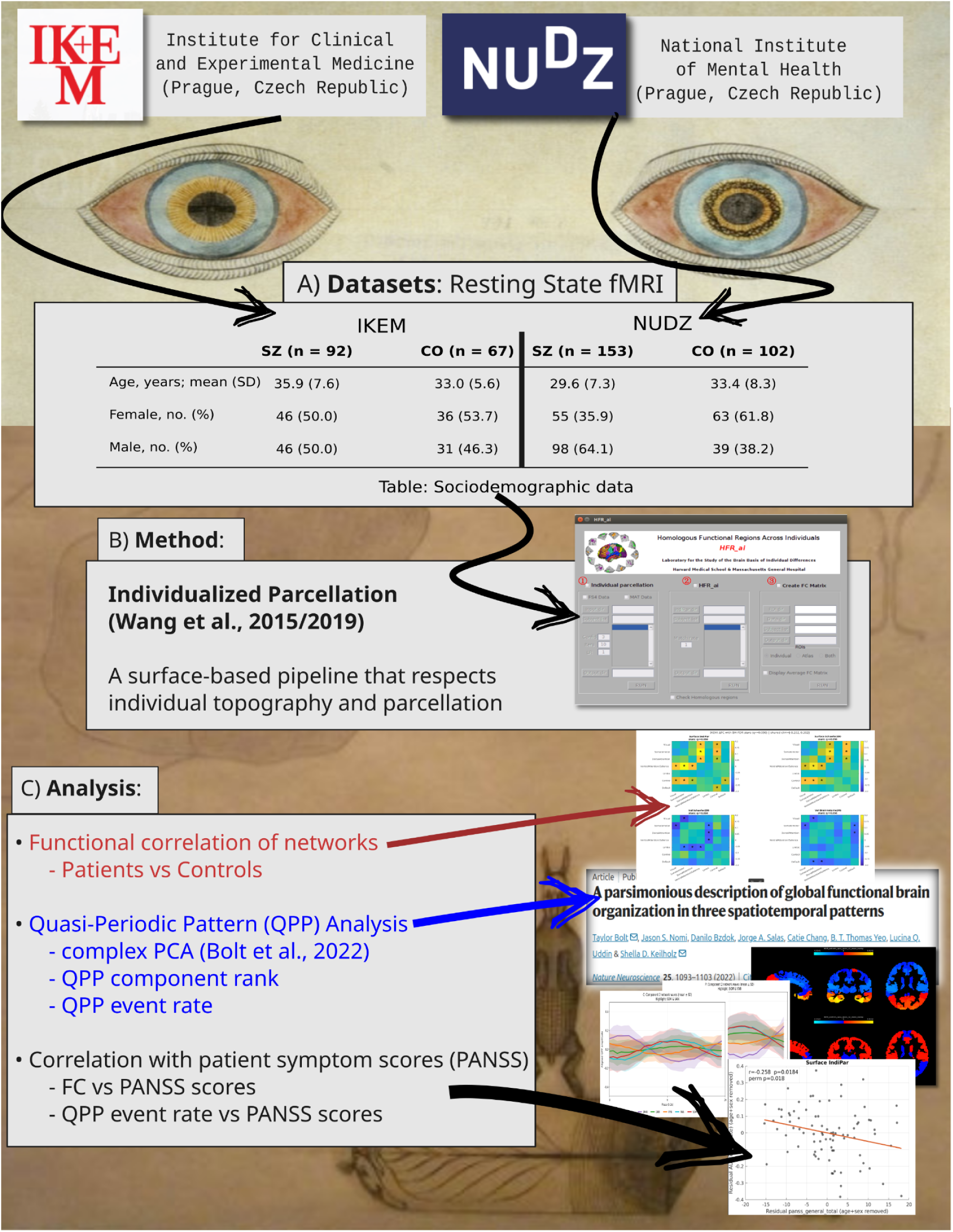
Study overview and analysis workflow. A) Age and Sex data for both datasets (IKEM and NUDZ), SZ = patients with schizophrenia spectrum disorder, CO = controls. B) Individualized surface-based parcellation (IndiPar), developed by Wang and colleagues, 2015/2019. Results from this method were compared with 3 other preprocessing/parcellation approaches, full description in methods section. C) Summary of analysis pipeline applied to all 4 preprocessing approaches: 1) Functional correlation (functional connectivity) between Yeo’s 7 networks for patients minus controls. 2) Detection of QPP component of task-negative/task-positive anticorrelation pattern using complex PCA toolbox, from Bolt and colleagues, 2022. Computation of QPP component rank, DMN-DAN opposition of QPP component, and QPP event rate. 3) Correlation of primary measures (delta FC and QPP event rate) vs patient symptom scores (PANSS general, positive, and negative). All analyses conducted on a subjectwise basis in parallel for both datasets and all 4 parcellation pipelines. Background Art: Paintings by August Natterer (1868-1933), a German artist on the Schizophrenia spectrum. Art is public domain and was accessed digitally at wikiart.org.

### 2.2 MRI Data Acquisition

MRI data were acquired on 3 Tesla Siemens scanners at both sites.

*IKEM*: Data were collected using a Siemens Trio scanner with a 12-channel head coil. Structural images were acquired using a T1-weighted MPRAGE sequence (TR = 2300 ms, TE = 4.6 ms, flip angle = 10°, voxel size = 1×1×1 mm). Functional images were acquired using gradient echo-planar imaging sensitive to the BOLD signal (TR = 2000 ms, TE = 30 ms, flip angle = 90°, voxel size = 3×3×3 mm, 30 slices, 400 volumes, ∼13 minute scan).

*NUDZ*: Data were collected using a Siemens Prisma scanner with a 64/20-channel head coil. Structural images were acquired using a T1-weighted MPRAGE sequence (TR = 2400 ms, TE = 2.3 ms, flip angle = 8°, voxel size = 0.7×0.7×0.7 mm). Functional images were acquired using gradient echo-planar imaging (TR = 2000 ms, TE = 30 ms, flip angle = 90°, voxel size = 3×3×3 mm, 30 slices, 300 volumes, 10 minute scan).

### 2.3 Resting-State Preprocessing

Resting-state fMRI data were preprocessed using the CONN functional connectivity toolbox (Whitfield-Gabrieli, S., and Nieto-Castanon, 2012) together with FreeSurfer surface reconstruction tools (Fischl, 2012). To account for scanner equilibration and initial signal instability, the first ten functional volumes were removed from each scan prior to analysis.

Functional preprocessing steps for all pipelines included slice-timing correction, motion correction via rigid-body realignment, segmentation of structural images into gray matter, white matter, and cerebrospinal fluid (CSF), and co-registration between functional and anatomical images. Subsequent denoising included regression of nuisance signals from white matter and CSF, motion parameters derived from realignment, and outlier regressors identified using artifact detection (ART). Temporal band-pass filtering was applied between 0.008 and 0.09 Hz to isolate low-frequency fluctuations relevant to resting-state functional connectivity.

For the two volumetric pipelines, preprocessing additionally included spatial normalization of functional and structural images to the MNI-152 template space using CONN default preprocessing settings. Global signal regression was not included.

For surface-based pipelines, preprocessing was performed in subject anatomical space without volumetric template normalization. Functional time series were projected from volumetric space to the cortical surface using FreeSurfer volume-to-surface mapping procedures and subsequently mapped to the fsaverage4 surface template. Fsaverage4 was chosen as the Wang IndiPar pipeline is optimized for it (Wang, 2019). The fsaverage4 representation contains 2,562 vertices per hemisphere (5,124 vertices total), providing a downsampled but topologically consistent surface representation suitable for subject-level connectivity analyses.

### 2.4 Parcellation Pipelines Overview

Four preprocessing and parcellation pipelines were compared, two surface-based approaches, and two volumetric CONN default approaches:

1. Individualized surface-based parcellation (IndiPar, Wang, 2015)
2. Surface-based Schaefer200 atlas (Schaefer, 2018)
3. Volumetric Schaefer200 atlas (Schaefer, 2018)
4. Volumetric Brainnetome246 atlas (Fan, 2016)

The Schaefer200 atlas was selected because it is widely used in large-scale functional connectivity studies and permits direct comparison between surface- and volume-based preprocessing approaches. The Brainnetome246 atlas was additionally included as an established connectivity-informed volumetric atlas to evaluate whether results generalized across different atlas definitions rather than reflecting idiosyncrasies of a single volumetric parcellation scheme. Surface-based pipelines operated on fsaverage4 surface time series, whereas volumetric pipelines extracted ROI time series after normalization to MNI-152 space (Mazziotta, 2001).

### 2.5 Individualized Surface-Based Parcellation (IndiPar)

Subject-specific cortical parcellations were generated using the individualized parcellation framework introduced by Wang and colleagues and implemented using the HFR_ai (homologous functional regions across individuals) toolbox (Wang et al., 2015; Wang et al., 2019). This approach estimates individualized functional network organization by combining a population-derived network template with each subject’s surface-based resting-state fMRI data.

In brief, the method begins with a population-level cortical network atlas derived from approximately 1000 healthy individuals. This template is iteratively refined for each subject using their resting-state functional connectivity patterns, producing individualized surface-based network maps that better reflect subject-specific cortical organization.

The resulting individualized networks are subsequently divided into spatially contiguous surface patches using FreeSurfer tools. These patches are then matched to homologous functional regions across subjects based on spatial overlap and geodesic distance along the cortical surface. Because individualized network organization varies between subjects, the resulting parcellations may differ in both the number and spatial extent of ROIs across individuals.

This variability was intentional and reflects inter-individual differences in cortical functional organization rather than forcing all subjects into a fixed atlas definition.

For subsequent analyses, individualized ROIs were assigned to canonical functional networks using the Yeo 17-network classification provided by the IndiPar outputs, which were then collapsed into the Yeo 7-network scheme to maintain comparability with atlas-based pipelines (Yeo et al., 2011). ROI time series were obtained by averaging vertex-level BOLD signals within each individualized patch.

Because all primary analyses were conducted at the subject level, variation in ROI number across participants did not impose methodological constraints on the functional connectivity or QPP analyses.

### 2.6 Group Atlas Parcellation Pipelines

For comparison to the IndiPar approach, three group-atlas pipelines were implemented.

In the surface-based Schaefer200 pipeline, functional time series in fsaverage4 surface space were extracted into the Schaefer200 atlas (pre-processing was identical to the IndiPar approach up to this point). Parcels were grouped according to the Yeo 7-network functional organization.

In the volumetric pipelines, ROI time series were extracted after normalization to MNI-152 space (i.e. CONN pre-processing defaults) using either the Schaefer200 volumetric atlas or the Brainnetome246 atlas. In each case, voxel time series within each atlas parcel were averaged to produce ROI-level signals.

### 2.7 Static Functional Connectivity

For each of the four pipelines, network-level time series were constructed by averaging signals from parcels assigned to each of the seven Yeo functional networks. Functional connectivity was then computed between the seven network time series, producing a 7 × 7 connectivity matrix (21 unique edges) for each subject. Pairwise Pearson correlations were computed between network time series for each subject and transformed using Fisher’s z-transformation.

Group differences in connectivity were tested using generalized linear models with group as the predictor and age and sex included as covariates where indicated. False discovery rate (FDR) correction was applied across all tested network edges and significant pairs represented as q-values less than 0.05.

### 2.8 Dynamic Analysis: Quasi-Periodic Patterns

Dynamic functional connectivity patterns were characterized using complex principal component analysis (cPCA) applied to ROI-level time series. This method was demonstrated by Bolt and colleagues (Bolt et al., 2022) to reliably identify the task-positive/task-negative anti-correlation pattern that is intrinsically recurring between Default Mode Network (DMN) and Dorsal Attention Network (DAN) regions of the brain. This pattern occurs on the infra-slow timescale (∼0.05–0.1 Hz) and has also been termed the quasi-periodic pattern (QPP) due to its semi-periodic recurrence (Majeed, 2011; Yousefi, 2018; Abbas, 2019; Maltbie, 2022; Daley, 2025; Watters, 2025). Consistent with the application of cPCA by Bolt and colleagues, 3 components were detected in the present study, as they found a significant elbow in explained variance after the top three components. One of the key findings from Bolt and colleagues, who applied cPCA on concatenated scans from Human Connectome Project data, was that the first component was always the Global-Signal (it explained the most variance of whole-brain intrinsic BOLD fluctuations); the QPP/DMN anticorrelation pattern was always the second component (it explained the second most variance).

As evidenced in the results section, the QPP is not always the second component when cPCA is run on subjectwise scans instead of concatenated data. To account for this in the present study, the QPP component was defined as the strongest non-global component among the top three cPCA components showing the strongest DMN-DAN opposition. Specifically, QPP-like components were selected from the top three subject-level cPCA components by excluding the component showing high similarity to a uniform whole-brain loading pattern (global_sim ≥ 0.75). Among the remaining candidates, the selected component was the one with the highest DMN-DAN composite score, defined from the absolute difference between mean Default and Dorsal Attention complex loadings and their phase offset.

Rank-1 prevalence was computed as the fraction of subjects for whom the selected non-global QPP-like component was identified as component 1. For DMN-DAN opposition and selected component explained variance, pairwise pipeline differences within each dataset were tested using two-sided Wilcoxon signed-rank tests, with BH-FDR correction across pairwise comparisons.

Because no established biological threshold defines a QPP “event”, a threshold-sweep approach was adopted. In this approach, QPP-like events were detected by thresholding the absolute time course of the selected QPP component. Amplitude thresholds were defined using control-derived quantiles ranging from 0.30 to 0.90 to capture a broad range of fluctuations in the component time course. At each threshold, contiguous above-threshold fluctuations in the component time series lasting between 6 and 30 seconds were classified as events, to permit individual heterogeneity in event duration. Event rate per minute was computed for each threshold, and the area under the threshold event rate curve was used as a summary measure of QPP event prevalence. The threshold-sweep approach was used to avoid reliance on a single arbitrary cutoff and to summarize event prevalence across a range of amplitudes. Control-derived thresholds were used so that event definitions reflected normative fluctuation levels rather than patient-specific variance.

### 2.9 Clinical Association Analyses

Associations between neuroimaging metrics (ΔFC in patients - controls, and QPP event rate) and clinical symptoms were tested within patient groups using partial correlations controlling for age, sex, and motion (framewise displacement) with two-sided Freedman-Lane permutation tests (10,000 permutations). Clinical measures included PANSS total, positive, negative, and general symptom scores.

For clinical association analyses, main-text heatmaps display effect sizes with nominal permutation significance (uncorrected), whereas BH-FDR-corrected family summaries across tested edges are reported in Supplementary Table S1.

### 2.10 Motion

Head motion decreases signal-to-noise ratio and is of great concern in structural and functional neuroimaging, especially for psychiatric patients who may move more in the scanner (Van Dijk, 2012; Makowski, 2018). To account for motion, motion parameters were regressed from the BOLD (blood oxygen level dependent) time series during preprocessing. However, because head motion can influence functional connectivity estimates (Satterthwaite, 2012) and patients in the present study exhibited greater mean framewise displacement than controls (supplemental figure 1), we performed additional analyses including mean framewise displacement as a covariate when conducting symptom-correlations.

### 2.11 Analysis Summary

All analyses were conducted independently for each dataset (IKEM and NUDZ) and for each preprocessing pipeline. The analysis strategy can therefore be described as mass-univariate. For each subject, ROI-level time series were extracted and used to compute both static functional connectivity (FC) in Yeo’s 7 networks and dynamic metrics derived from quasi-periodic patterns (QPPs).

Clinical association analyses were conducted within patient groups (IKEM and NUDZ) to evaluate relationships between FC and PANSS symptom scores (general, positive, and negative).

## 3. Results

### 3.1 Patient-Control Differences in Static Functional Connectivity

Individualized surface-based parcellation revealed the strongest (highest magnitude of change) patient-control differences in static functional connectivity (ΔFC). Using the Yeo-7 network framework, the IndiPar pipeline identified the largest number of significant FC edges (two-sided Welch t-tests, BH-FDR correction within each dataset × pipeline across the 21 unique Yeo-7 edges, q ≤ 0.05). These differences were distributed across multiple network pairs, with prominent effects involving default mode network (DMN), dorsal attention network (DAN), and frontoparietal control network (FPN) interactions (Figure 2).

**Figure 2:**
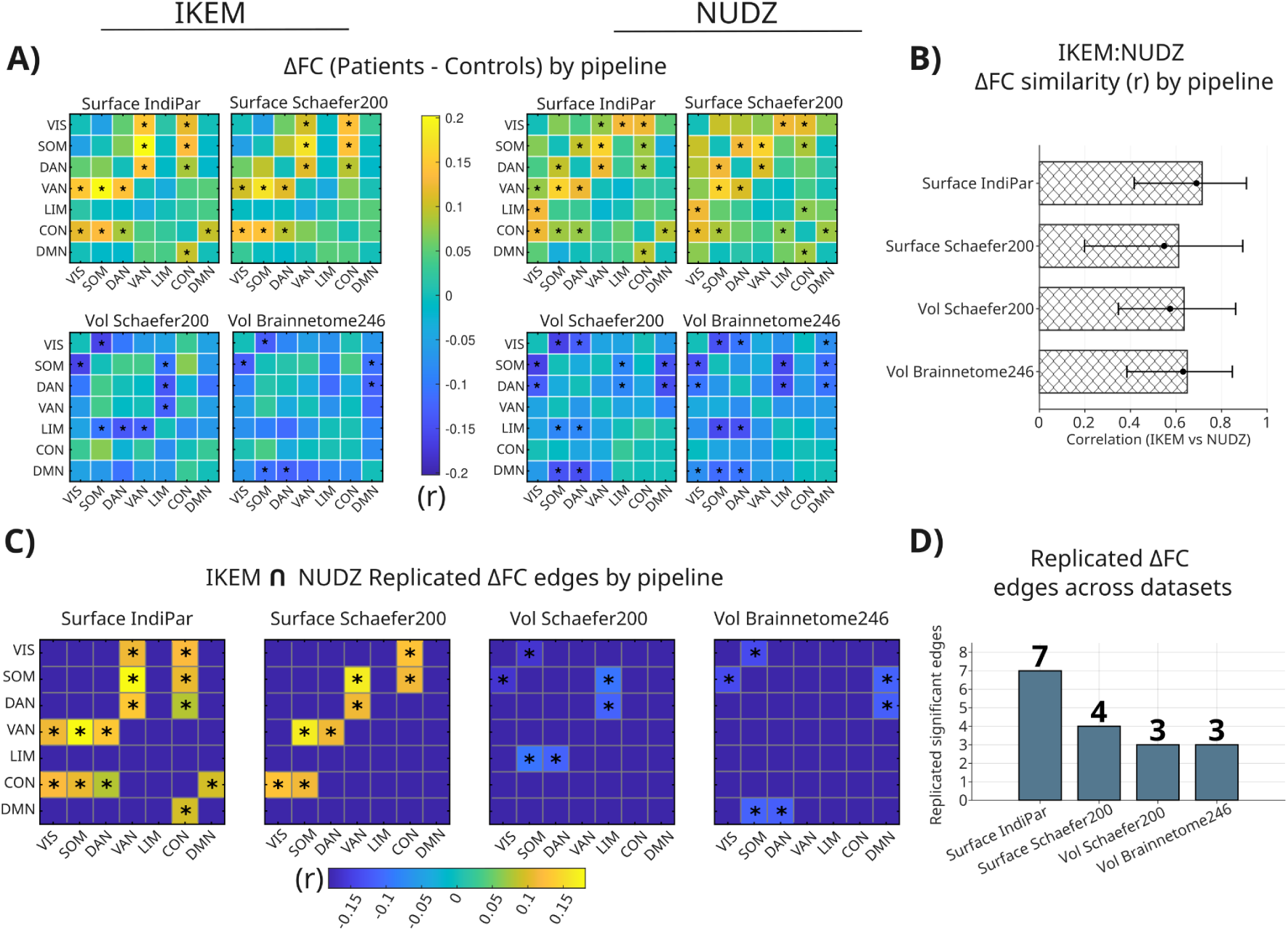
Functional connectivity (Patients minus controls). A) Differences in functional correlation between Yeo’s 7 networks in the IKEM and NUDZ datasets, for each of the 4 parcellation pipelines. Stars = significant edges. Edges were considered significant at q ≤ 0.05. (Two-sided Welch’s t-tests with BH-FDR correction across edges (m = 21). B) Similarity (correlation) between datasets by pipeline. IndiPar trends higher but does not reach significance. C) Summary heatmaps showing overlap of significant edges between the two datasets. Note that surface-based pipelines have much greater magnitude (color intensity) of ΔFC and are positive (signifying patients > controls for those network pairs), while the volumetric pipelines show weaker ΔFC and little overlap for network pairs and ΔFC direction with surface pipelines. D) Histogram counts of cross-dataset replicated edges.

Surface-based Schaefer200 parcellation detected a smaller subset of significant ΔFC edges but generally reproduced the spatial pattern observed with IndiPar. In contrast, volumetric pipelines showed smaller effect sizes and fewer significant pairs. The Brainnetome246 pipeline consistently produced the lowest number of significant ΔFC edges.

Importantly, several edges detected with IndiPar were not identified using atlas-based approaches, indicating that fixed parcellation schemes may obscure subject-specific connectivity alterations. These patterns were consistent across IKEM and NUDZ cohorts, suggesting that individualized surface-based parcellation better preserved patient-control differences in large-scale network connectivity.

### 3.2 Cross-Dataset Consistency of FC

To evaluate reproducibility across independent cohorts, ΔFC matrices derived from IKEM and NUDZ were compared directly. Surface-based pipelines trended higher for cross-dataset similarity than volumetric pipelines, with the IndiPar pipeline showing the highest mean correlation between datasets. Replicated effects were defined as edges that survived BH-FDR correction in both IKEM and NUDZ (Figure 2, C-D).

Although the difference in cross-dataset similarity between pipelines did not reach statistical significance, the consistent trend toward higher reproducibility in surface space suggests that individualized parcellation may better preserve biologically meaningful connectivity structure across datasets collected at different sites. IndiPar also yielded a higher number of replicated ΔFC edges (7) between IKEM and NUDZ compared to the other pipelines (Figure 2, C).

### 3.3 Differences in QPP Component Characteristics Between Pipelines

Dynamic analyses using complex principal components analysis (cPCA) revealed systematic differences in quasi-periodic pattern (QPP) characteristics across preprocessing pipelines (Figure 3). Surface-based pipelines, particularly IndiPar, more frequently identified canonical QPP components characterized by strong anti-correlation between DMN and DAN networks.

**Figure 3:**
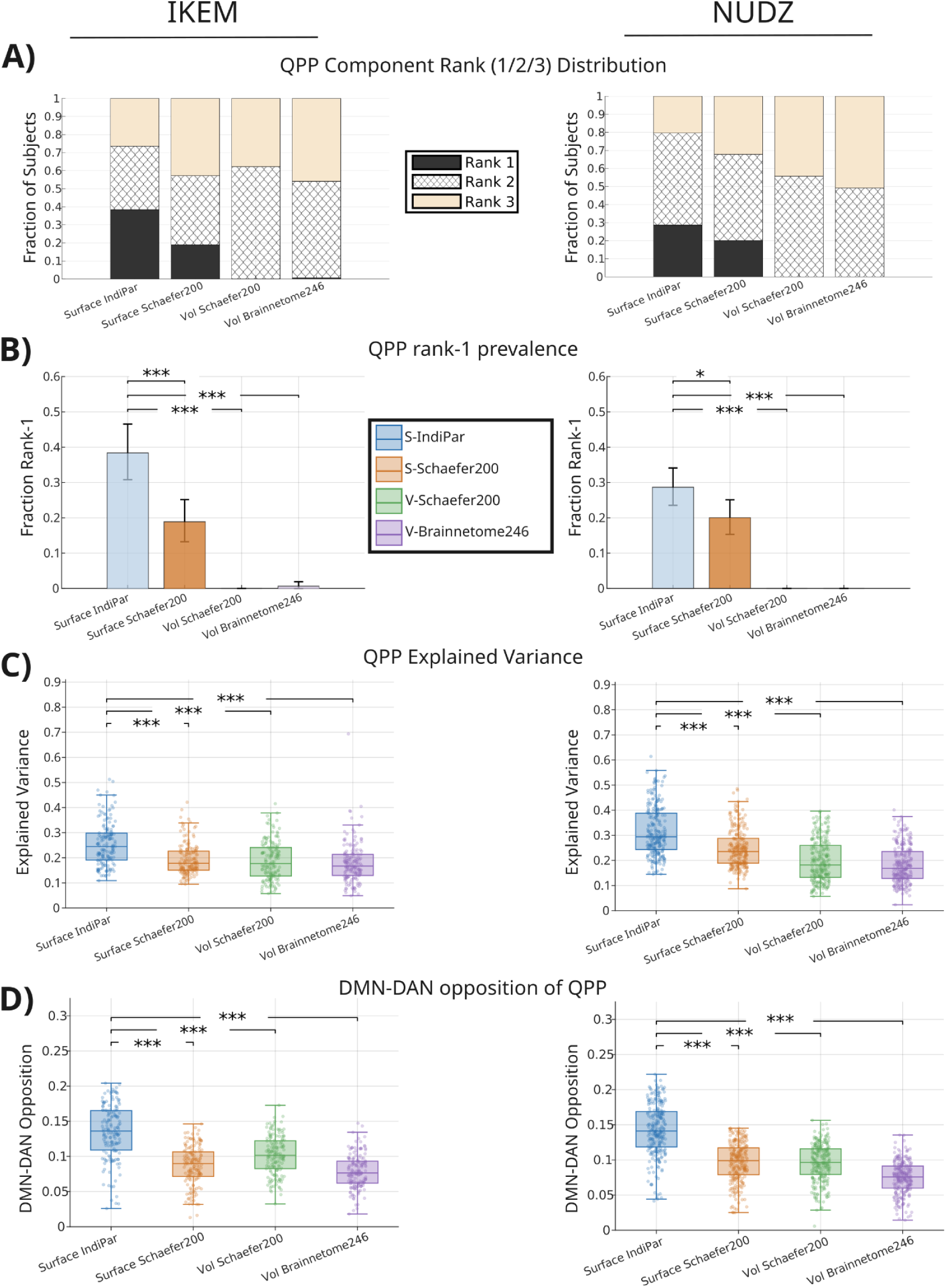
Detection of QPP dynamics across preprocessing pipelines. From top to bottom rows, outcomes are compared across both datasets using the merged patient and control cohorts (IKEM and NUDZ) and the four preprocessing pipelines for **A)** distribution of the cPCA component rank (1–3) assigned to the QPP, **B)** prevalence of rank-1 QPP components, **C)** variance explained by the selected QPP component, and **D)** degree of DMN-DAN opposition for the selected QPP component. Significance brackets indicate pairwise pipeline comparisons with BH-FDR correction within dataset. * q < 0.05, ** q < 0.01, *** q < 0.001. The individualized surface-based pipeline (IndiPar) produced a significantly higher prevalence of rank-1 QPP components (shown as fraction ± bootstrap 95% confidence intervals; 5,000 resamples; McNemar tests with BH-FDR correction), higher explained variance of the selected component (two-sided Wilcoxon signed-rank tests; all pairwise comparisons vs IndiPar q < 0.001 after BH-FDR correction within dataset), and stronger DMN-DAN opposition (two-sided Wilcoxon signed-rank tests; all pairwise comparisons vs IndiPar q < 0.001 after BH-FDR correction within dataset) relative to atlas-based pipelines.

QPP-like components were still detected in volumetric pipelines; however, these components were less frequently identified as the dominant (rank-1) component and generally exhibited weaker DMN-DAN opposition and lower explained variance. Instead, volumetric pipelines more frequently produced dominant components resembling global-signal fluctuations, consistent with prior observations that global signal tends to explain the greatest variance in cPCA decompositions (Bolt, 2022). These findings suggest that standard volumetric preprocessing may reduce the prominence of canonical association-network QPP dynamics relative to global fluctuations.

Across both datasets, the IndiPar pipeline showed a substantially higher prevalence of QPP components identified as rank-1 compared with atlas-based pipelines (Fig. 3; IKEM and NUDZ; chi-square tests of homogeneity comparing rank-1 vs non-rank-1 selections; BH-FDR corrected q values ≤ 0.023 across IndiPar-versus-pipeline comparisons). In addition, the QPP components selected by the IndiPar pipeline explained significantly greater variance in the complex-PCA decomposition than those identified by atlas-based pipelines (two-sided Wilcoxon signed-rank tests; all pairwise comparisons vs IndiPar significant after BH-FDR correction across pipelines within dataset, q < 0.0001). Finally, the selected QPP components exhibited significantly stronger Default Mode Network-Dorsal Attention Network (DMN-DAN) opposition in the IndiPar pipeline relative to the comparison pipelines in both datasets (two-sided Wilcoxon signed-rank tests; all pairwise comparisons vs IndiPar significant after BH-FDR correction across pipelines within dataset, q < 0.0001). Together, these results indicate that individualized parcellation preferentially identifies dominant QPP components that capture greater variance and exhibit stronger canonical DMN-DAN antagonism. In other words, individualized parcellation appears to improve detection of QPP dynamics.

**Figure 4:**
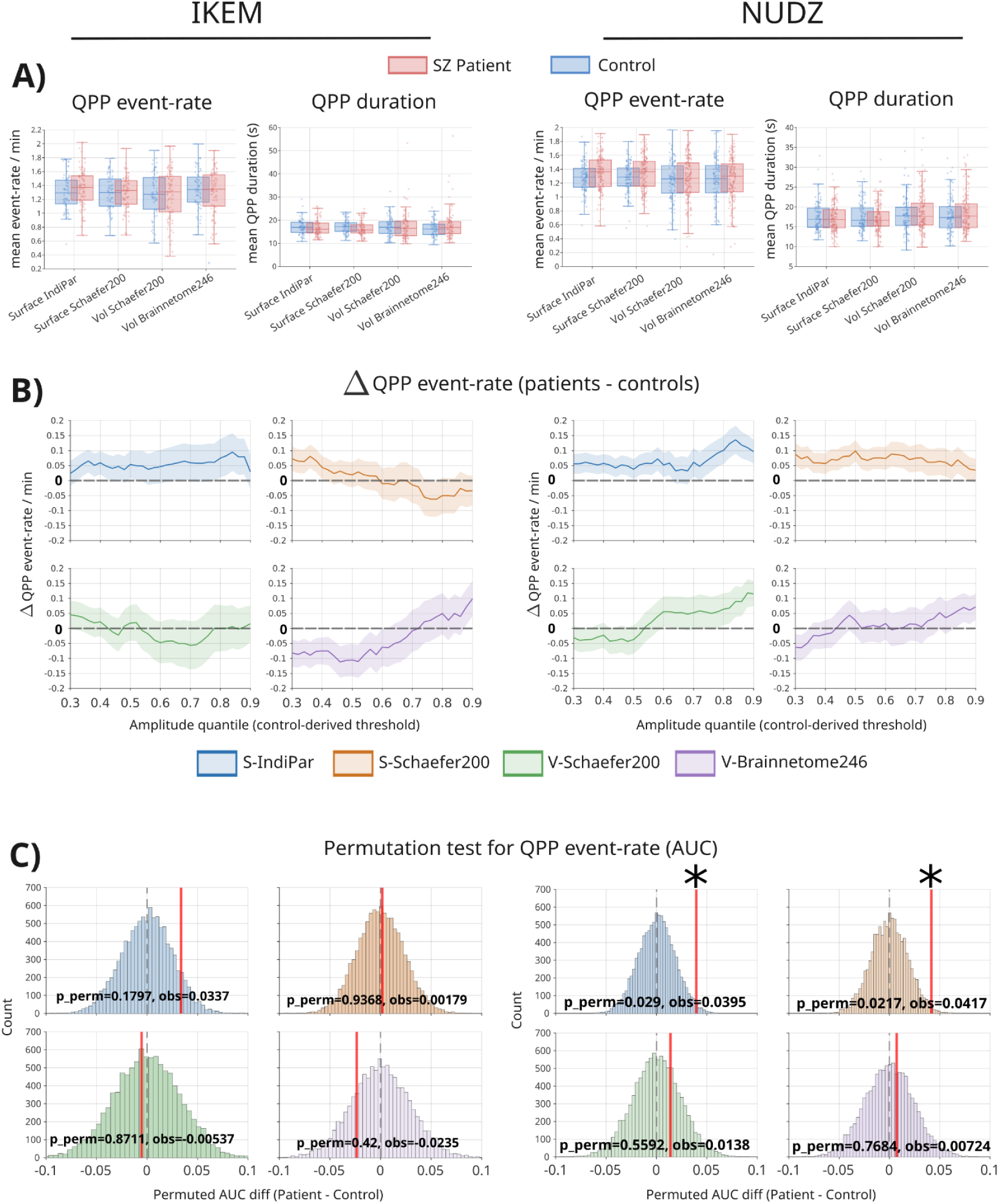
Comparison of QPP duration and event rate per minute in SZ patients vs controls (Left: NUDZ, Right: IKEM). **A)** Descriptive boxplots show subject-level mean QPP event rate and mean QPP duration (averaged across thresholds) for patients and controls in each pipeline. Group differences in panel A were evaluated using Wilcoxon rank-sum tests within each dataset and pipeline. Event rate distributions showed a modest trend toward higher values in patients under the individualized surface pipeline, whereas QPP duration did not differ consistently between groups across pipelines. **B)** Descriptive mean Δ event rate curves (patients minus controls) across amplitude thresholds show that the individualized surface pipeline (IndiPar) produced a positive patient-control difference across most thresholds in both datasets, while atlas-based pipelines showed smaller, inconsistent, or directionally reversed effects. **C)** To summarize threshold-sweep differences, a subject-level area-under-the-curve (AUC) metric was computed for each subject and compared between groups using two-sided permutation tests (10,000 permutations). The patient-control AUC difference showed a consistent patient > control trend for the individualized pipeline across datasets, reaching statistical significance in NUDZ for IndiPar and Surface-Schaefer200, while other pipelines were not significant.

### 3.4 Patient-Control Differences in QPP Event Rate

QPP event rate was quantified across amplitude thresholds and summarized using area-under-the-curve (AUC) metrics. Across both cohorts, the IndiPar pipeline trended higher for QPP event rates in patients relative to controls. The patient-control difference in QPP event rate AUC was significant in NUDZ for IndiPar and Surface-Schaefer200 (two-sided permutation tests, 10,000 permutations, Figure 3. C-D). The patient-control difference was similar in IKEM, but with significantly fewer subjects the difference did not reach significance after 10,000 permutations (Figure 3, D).

Surface-based Schaefer200 parcellation also detected patient-control differences across thresholds, but the direction of change was not consistent between NUDZ and IKEM as it was in the IndiPar pipeline. Volumetric pipelines produced the weakest and least consistent differences in event rate across thresholds, consistent with reduced sensitivity to dynamic network activity under atlas-based volumetric preprocessing (Figure 3, C-D).

### 3.5 QPP Duration

No qualitative differences in the average QPP duration were observed between patients and controls and thus the QPP duration was not further analyzed. However, the subjectwise results of the QPPs identified with cPCA in the present study are worth contrasting with reported QPP durations obtained in concatenated group-level data. Bolt and colleagues for example, found in resting state HCP data that QPPs tend to last ∼20 seconds per cycle (one full wave from zero to 2***π***). When detecting QPPs on the subjectwise level (Figure 3, A) it is apparent that QPP duration (and thus event rate) vary substantially between subjects, with averages in the ∼15 second range and variances covering anywhere from roughly 10-30 seconds. In some sliding-window based detection methods (Xu, 2025), the user defines a window length which effectively forces the algorithm to converge on an average pattern of ∼20 seconds. Subjectwise detection of QPPs suggests group-detection may be a problematic approach, as many subjects simply do not show QPPs of that length in resting state data, especially over short scan lengths.

### 3.6 Clinical Associations: QPP Event Rate

Associations between QPP event rate and symptom severity were evaluated using PANSS total and subscale scores while controlling for age, sex, and motion. Under the IndiPar pipeline, QPP event rate demonstrated significant correlations with multiple PANSS measures in the IKEM cohort but not the NUDZ cohort (Pearson partial correlations controlling for age, sex, and mean FD; two-sided Freedman-Lane permutation tests, 10,000 permutations, Figure 5, B). These associations should be considered exploratory. Although nominal permutation-significant PANSS associations were observed in IKEM under the IndiPar pipeline, they did not show cross-dataset replication, and none survived full correction of age, sex, motion plus FDR, again demonstrating the precarious nature of psychiatric fMRI-biomarker discovery. The discrepancy between datasets may partly reflect cohort and site-related differences, as weaker effect magnitudes in the NUDZ cohort have also been reported in an independent self-agency fMRI study using the same dataset (Volfíková et al., 2025).

**Figure 5:**
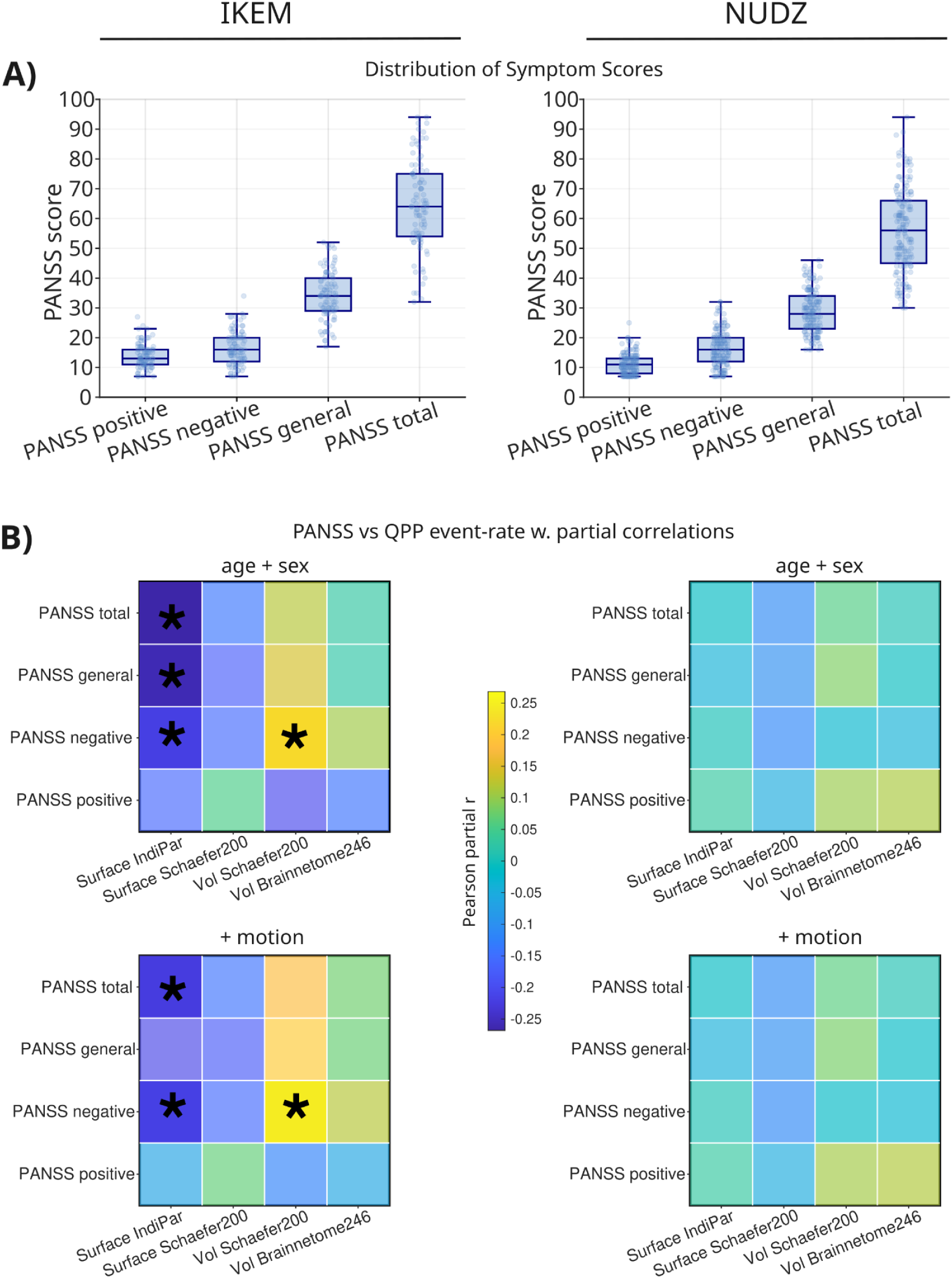
Relationship between QPP event rate and symptom scores. **A)** Distribution of symptom scores. Note qualitative similarity of symptom ranges across datasets. **B)** Heatmaps show partial correlations between QPP event rate area under the curve (AUC) and PANSS symptom domains in patients with schizophrenia. Analyses were performed separately for the IKEM and NUDZ datasets using four preprocessing pipelines. Correlations were computed using Pearson partial correlation while controlling for age and sex (B, top row) as well as age, sex, and motion (B, bottom row). Cell color indicates the correlation coefficient (r), with a shared color scale across panels. Asterisks denote nominal permutation significance (two-sided Freedman-Lane tests, 10,000 permutations, p < 0.05, uncorrected); corrected summaries are reported in Supplementary tables. QPP event rate AUC summarizes the frequency of detected QPP events across the threshold sweep for each subject. Note the largely unchanged relationship between QPP event rate and PANSS for either dataset after accounting for increased patient motion as a covariate.

Significant associations were largely absent when using Schaefer200 surface or volumetric pipelines. Partial correlations were reduced toward zero and generally failed to reach significance under atlas-based approaches in IKEM. These findings suggest that individualized surface-based parcellation preserved more nominal brain-symptom signal in IKEM, although these associations were exploratory and did not replicate across datasets. Additionally, the results indicate that (across pipelines) QPP event rate is not heavily affected by motion as a covariate. The lack of cross-dataset replication between IKEM and NUDZ urges caution in QPP trait level inference related to SZ symptomatology.

### 3.7 Clinical Associations: Functional Connectivity

Edge-wise correlations between static FC and PANSS symptom scores were mixed, showing the strongest positive correlations under the IndiPar pipeline and negative correlations in volumetric pipelines. Several FC edges demonstrated significant associations with PANSS total and subscale scores after motion correction (Figure 6, A). Many of these associations involved interactions between DMN, attention networks, and control networks.

**Figure 6:**
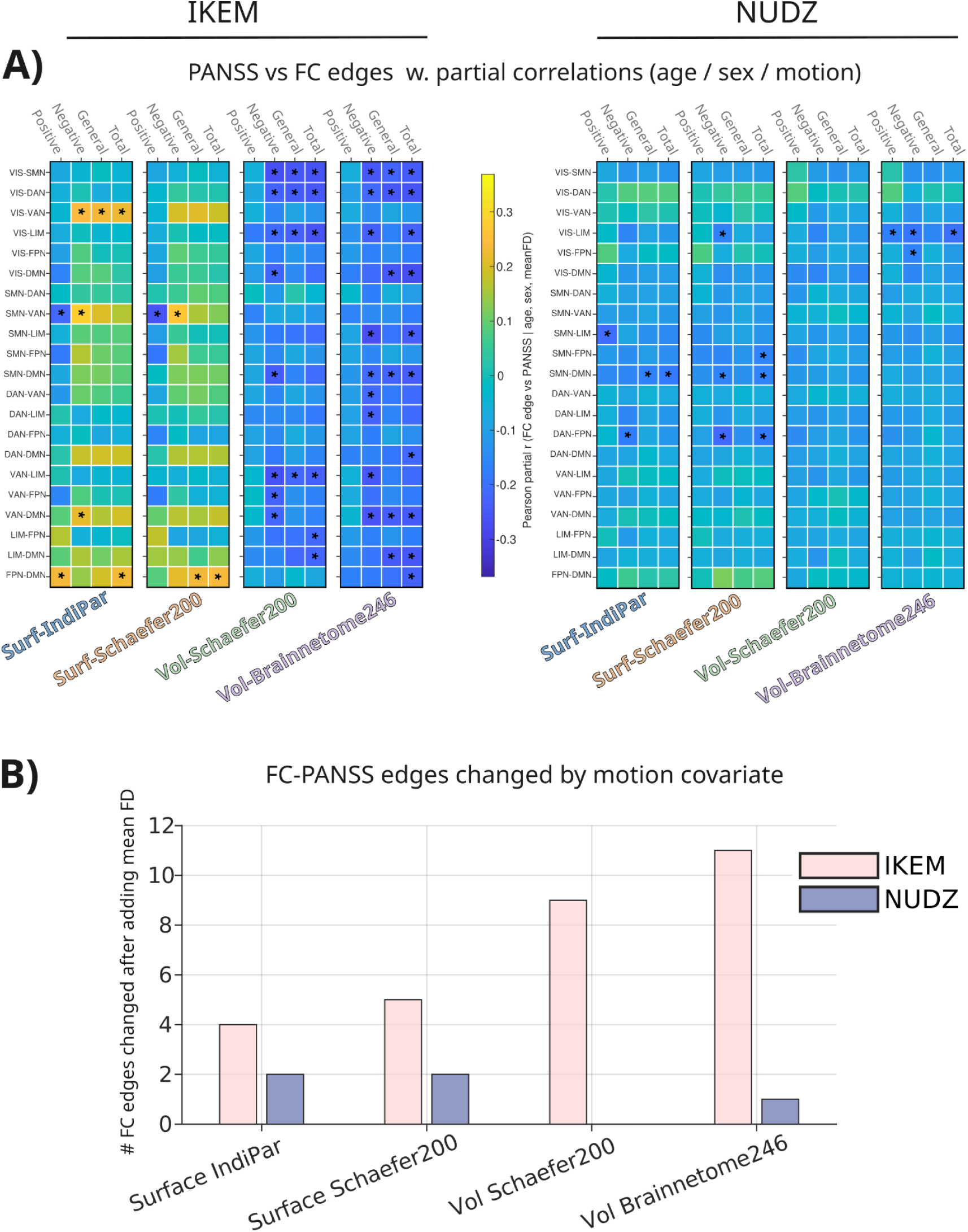
Relationship between FC, symptom scores, and motion. A) Heatmaps showing partial correlations between patient functional connectivity and PANSS symptom domains. Each column represents a network-pair connectivity edge, and each row represents a PANSS domain. Cell color indicates the correlation coefficient (r), with a shared color scale across panels. Significance was defined as nominal permutation p < 0.05 (two-sided Freedman-Lane tests, 10,000 permutations, uncorrected). Non-significant cells are slightly faded for visualization. B) Summary of changed FC-PANSS edges after inclusion of motion covariate. A high number of FC-PANSS significant edges were altered by inclusion of motion as a covariate, even after motion had already been regressed out as a confound during preprocessing, consistent with the understanding that functional correlations are strongly influenced by motion. In the PANSS vs QPP event rate analysis (figure 5) only a single significant edge was changed across all pipelines. See supplemental for pre-motion FC-PANSS plots. Note there is little agreement between the two datasets for FC vs PANSS relationship.

Surface-based Schaefer200 parcellation detected a smaller subset of these associations. Before motion was included as a covariant, volumetric pipelines detected only a limited number of significant FC-PANSS relationships (Supplemental). Notably, the direction and magnitude of several correlations varied substantially across pipelines, underscoring the sensitivity of brain-behavior relationships to preprocessing and parcellation choices. Comparing the pipeline results side-by-side does not reveal any consistent FC edges vs PANSS symptom scores, highlighting the pitfall potential of single-pipeline + single-dataset approaches to psychiatric biomarker discovery.

### 3.8 Motion as a covariate

After including motion as a covariate, the overall pattern of QPP event rate vs PANSS correlations was largely unchanged (only 1 significant edge changed, figure 5, B). This is consistent with previous work showing that motion does not significantly contribute to QPPs (Yousefi, 2018). In contrast, numerous FC-PANSS associations, particularly in volumetric pipelines, were sensitive to motion covariation (two-sided t-tests followed by BH-FDR correction within each dataset × pipeline across 21 edges, results after motion in Figure 6). These results match an understanding that resting state functional connectivity results are very susceptible to motion effects (Kopal et al., 2020; Maziero et al., 2020). Notably, while there is some overlap in significant FC-PANSS edges in surface-surface and volume-volume parcellations *within* each dataset, there is little overlap in FC vs symptom scores *between* the datasets. No clear cross-dataset FC vs symptom pairs emerged across datasets and pipelines, again underscoring the tenuous nature of single-dataset schizophrenia neuroimaging findings based on univariate features. This sensitivity of FC-symptom associations to preprocessing and analytical choices is consistent with recent work demonstrating substantial shifts in observed schizophrenia connectivity effects across denoising strategies (Tomecek et al., 2026). Figure 6 shows FC vs PANSS results after motion was included as a covariate, see Supplementary for before motion results.

## 4. Discussion

Across two independent schizophrenia cohorts, individualized surface-based parcellation consistently enhanced detection of canonical quasi-periodic pattern (QPP) dynamics relative to standard atlas-based approaches. IndiPar produced stronger DMN-DAN opposition, higher explained variance of canonical QPP components, and greater prevalence of rank-1 QPPs, suggesting improved sensitivity to large-scale intrinsic brain dynamics. These effects were not explained solely by surface-based preprocessing, as the surface-based Schaefer200 pipeline did not reproduce the same magnitude of enhancement. Cross-dataset similarity of schizophrenia-related FC effects showed partial improvement under IndiPar, whereas symptom-related associations remained substantially variable across datasets, consistent with the known heterogeneity and limited reproducibility of brain-symptom correlations in psychiatric neuroimaging.

### 4.1 Revisiting Hypotheses and Summary

The prediction that IndiPar would more sensitively detect infraslow dynamics such as QPPs was strongly supported by the results (Figure 3). IndiPar consistently produced higher QPP rank, stronger DMN-DAN opposition, and greater explained variance of canonical QPP components relative to atlas-based approaches, suggesting improved sensitivity to large-scale intrinsic dynamics. Importantly, these effects could not be attributed solely to surface-based preprocessing, as the surface-based Schaefer200 pipeline used identical surface preprocessing but did not reproduce the same magnitude of effects. This suggests that individualized parcellation itself contributes to improved detection of intrinsic brain dynamics beyond surface representation alone.

The hypothesis that individualized parcellation would improve similarity of results between independently acquired datasets received partial support. IndiPar produced more similar cross-dataset FC differences between patients and controls (Figure 2), suggesting that individualized approaches may reduce some sources of variability associated with standard atlas-based normalization. However, this increased consistency did not clearly extend to symptom-related FC associations or QPP prevalence measures across datasets. These findings align with previous reports that schizophrenia neuroimaging effects often show limited cross-dataset generalizability, potentially reflecting substantial biological heterogeneity, site effects, and methodological sensitivity within diagnostic categories (Wolfers, 2018; Voineskos, 2024).

Lastly, we hypothesized that the IndiPar approach would preserve stronger associations between functional imaging measures and patient symptom severity (PANSS). Results in this domain were more variable and dataset dependent, which is unsurprising given that symptom-severity correlations represent higher-order associations influenced not only by neurobiological variation, but also by the reliability and heterogeneity of clinical symptom assessment. In the IKEM dataset, IndiPar preserved a greater number of significant QPP vs. symptom correlations before and after motion correction. However, these effects were substantially weaker and did not reach significance in the NUDZ dataset, consistent with prior observations of attenuated schizophrenia-related effect sizes in this sample (Volfíková et al., 2025).

Interestingly, QPP event rate was negatively correlated with PANSS symptom domains (general, positive, and negative symptoms) (Figure 3B). Given that in both datasets QPP event rate either trended higher or was significantly elevated in patients relative to controls, it was initially expected that QPP event rate might positively correlate with symptom severity, particularly given evidence that QPPs contribute more strongly to functional connectivity in schizophrenia (Briend, 2020). However, all three symptom domains instead demonstrated inverse relationships with QPP event rate. One possible interpretation is that the anti-correlated relationship between QPP event rate and symptoms reflects compensatory or adaptive network dynamics in some individuals, or alternatively non-linear relationships between intrinsic network activity and symptom burden. However, these interpretations remain speculative and will require replication in larger cohorts.

Interpretation of dynamic functional connectivity measures remains an active methodological area in neuroimaging (Laumann et al., 2024). Unlike sliding-window correlation approaches, the quasi-periodic patterns examined here reflect structured spatiotemporal activity patterns identified using complex principal component analysis (Bolt et al., 2022). QPPs have been observed across multiple datasets and analysis approaches (Majeed, 2011; Thompson, 2014; Yousefi, 2018; Abbas, 2019a), suggesting that they represent reproducible features of intrinsic brain activity rather than statistical artifacts of stationary covariance structure. In the present study, QPP components were selected after excluding the dominant global component identified by cPCA, minimizing potential contributions from global signal fluctuations.

The primary contribution of this study is methodological rather than clinical. Our findings demonstrate that preprocessing and parcellation choices substantially influence detection of intrinsic brain dynamics, emphasizing the importance of evaluating analytic pipelines when interpreting resting-state neuroimaging results, especially in single-dataset + single-pipeline studies.

### 4.2 Consideration of biomarker discovery for psychiatric categories

Clinical associations between QPP dynamics or functional connectivity and PANSS scores were not robust across cohorts, preprocessing pipelines, multiple-comparison corrections and motion covariate control. While nominal associations were observed in specific dataset-pipeline combinations, these effects did not replicate across datasets and were sensitive to the inclusion of motion as a covariate.

These findings underscore the fragility of cross-sectional brain-symptom relationships in resting-state fMRI and highlight the importance of evaluating reproducibility across datasets and pipelines. Rather than providing evidence for stable biomarkers, the present results suggest that apparent associations in rs-fMRI may reflect dataset-specific variance, preprocessing-dependent effects, or residual confounding. This reinforces the need for multi-dataset validation, standardized analytic frameworks, and cautious interpretation of single-cohort findings in psychiatric neuroimaging. This issue is not specific to SZ, as the search for reliable psychiatric biomarkers in domains such as depression and post-traumatic stress disorder suffers from similar heterogeneity issues (Wen et al., 2025). Additionally, the results contribute to a growing body of work suggesting that designing fMRI studies as A/B comparisons based on legacy diagnostic definitions may limit progress in understanding psychiatric disorders (Hengartner, 2017).

### 4.3 Limitations

Several limitations should be considered when interpreting these results.

First, the two datasets were acquired on different MRI scanners with slightly different sequences, including e.g. structural imaging resolutions. Although such differences may contribute to cross-dataset variability despite identical preprocessing pipelines, the use of independent cohorts acquired under non-identical conditions also represents a strength of the study, as it provides a more realistic test of methodological robustness and cross-site generalizability than would be possible in a single-site dataset.

Second, scan length is an important consideration for dynamic analyses, as QPP events occur relatively infrequently in resting-state data (Laumann, 2024). Given the relatively short resting-state acquisitions analyzed (∼10–13 minutes), the number of observable QPP events per subject was limited, and longer acquisitions would likely provide more stable estimates of dynamic network properties. However, scan durations of this length are typical of many clinical resting-state studies, particularly in psychiatric populations where participant burden and motion become increasingly important considerations.

Third, individualized parcellation produces variable numbers and spatial extents of cortical parcels across subjects. Although analyses in the present study were conducted at the network level (Yeo’s 7) to maintain comparability across pipelines, variability in parcel counts may influence fine-scale connectivity estimates relative to fixed atlas approaches.

Fourth, the schizophrenia cohorts analyzed here likely contain substantial clinical heterogeneity. Such heterogeneity may contribute to the limited replication of symptom associations across datasets and complicates interpretation of brain-behavior relationships.

Another methodological consideration concerns the role of global signal regression (GSR). Because GSR can increase the prominence of DMN-DAN anticorrelation patterns in resting-state BOLD data (Yousefi et al., 2018), future work should examine how individualized parcellation interacts with preprocessing pipelines that include GSR to determine whether the present findings generalize across denoising strategies.

Finally, interpretation of dynamic functional connectivity measures remains an area of active methodological discussion (Laumann, 2024). Although complex PCA provides a principled approach to identifying recurring spatiotemporal patterns (Bolt, 2022), the biological interpretation of dynamic components such as QPPs remains an open question in the field.

## 5. Conclusions and Future Directions

The present study demonstrates that preprocessing and parcellation choices can substantially influence both static and dynamic resting-state fMRI findings in schizophrenia. Across two independent cohorts, individualized surface-based parcellation consistently increased sensitivity to quasi-periodic patterns (QPPs), yielding higher QPP component rank prevalence, stronger canonical DMN-DAN opposition, and greater explained variance relative to standard atlas-based pipelines. IndiPar also detected stronger patient-control differences in static functional connectivity and preserved more symptom associations in one of the two datasets (IKEM). Together, these results suggest that individualized surface-based approaches may better capture subject-specific network dynamics that are partially obscured when fixed atlas frameworks are imposed.

At the same time, the limited replication of symptom associations across the two datasets highlights the persistent challenge of heterogeneity in psychiatric neuroimaging. Even when analytic pipelines were standardized, brain-behavior relationships showed little cross-dataset agreement, consistent with growing evidence that categorical diagnostic groupings may aggregate individuals with diverse underlying neural mechanisms (Wolfers, 2018; Voineskos, 2024). Rather than identifying single biomarkers for schizophrenia, approaches that emphasize individualized brain organization and subject-level variability may provide a more productive framework for studying psychiatric illness - one that embraces variability over category. Future work combining individualized parcellation methods with longer resting-state acquisitions, naturalistic tasks (Gal, 2022), multisite datasets, and dimensional clinical measures may help clarify how intrinsic brain dynamics relate to symptom variability and cognitive function across psychiatric populations.

## Code and Data Availability

Complex principal component analysis was performed using the publicly available implementation released by Bolt and colleagues for fMRI analyses (Bolt, 2022). Individualized surface parcellation was performed using the individualized parcellation / homologous functional regions framework described by Wang and colleagues (Wang, 2019), together with the publicly distributed HFR/LiuLab tools. All additional MATLAB code used for dataset-specific preprocessing orchestration, quality-control summaries, statistical analyses, and figure generation was developed in-house for the present study and is available from the corresponding author upon reasonable request. Primary code examples are available on GitHub at: https://github.com/Dalimear/Schizophrenia-qpp-cPCA-dynamics. Owing to data-use and privacy restrictions applying to the schizophrenia datasets analyzed here, the underlying imaging and clinical data are not publicly posted but may be made available through the relevant institutional data access procedures where permitted.

## Acknowledgments

This work was Supported by ERDF-Project No. CZ.02.01.01/00/23_020/0008560, part of BRAINSCAPE and the Czech Academy of Sciences, Institute of Computer Science (https://cobra.cs.cas.cz/). Funding was provided through the European Union and Czech Academy of Sciences. The authors acknowledge the use of ChatGPT (GPT-5.5, OpenAI) for improving grammar and language of the late-stage manuscript. All literature review, bibliography construction, citation, and initial drafting was done manually.

## Declaration of Competing Interests

The authors have no competing interests to disclose.

## Notes

https://github.com/Dalimear/Schizophrenia-qpp-cPCA-dynamics

### Competing Interest Statement

The authors have declared no competing interest.

