## Supplementary figs and table for "Individualized surface parcellation enhances characterization of resting-state brain dynamics and their alterations in schizophrenia"

**Supplemental**:

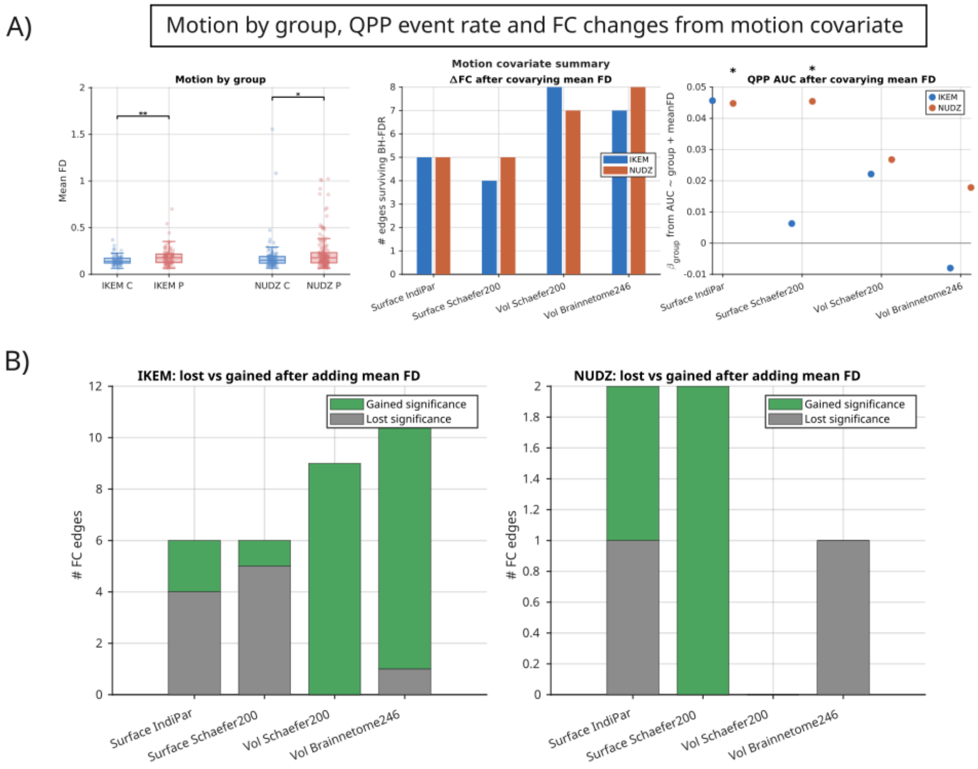


**Figure S1**: Motion by dataset, and QPP event rate and FC changes from motion covariate

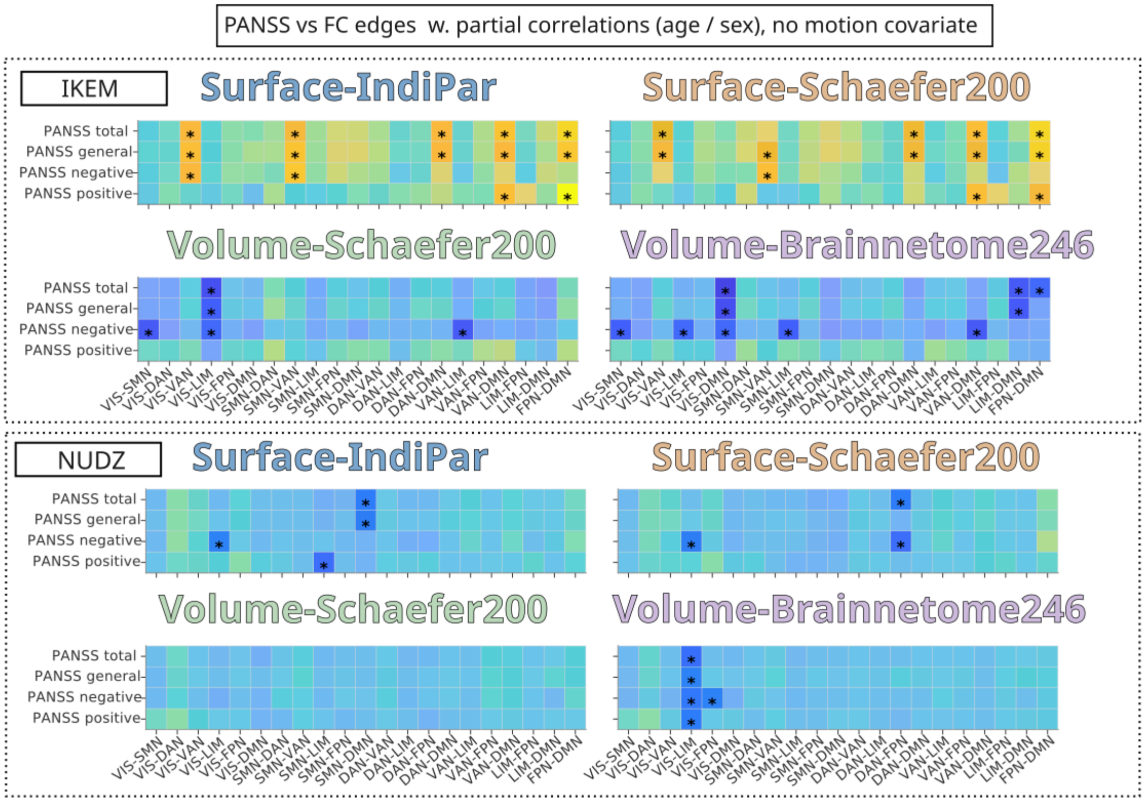


**Figure S2**: FC vs PANSS without motion covariate.

Supplemental Table S1: Statistical Comparison Tests and Corrections. Green = significant q/p

### Supplementary Table S1 (Page 1 of 4)

| analysis | dataset | pipeline | measure | family_size | test_and_correction | threshold | n_significant | min_p_or_q |
|---|---|---|---|---|---|---|---|---|
| Delta FC (Patient - Control) | IKEM | Surface IndiPar | Yeo7 between-network FC (z); 21 edges | 21 | Welch t-test (edgewise) + BH-FDR within dataset×pipeline across 21 edges | q<=0.05 | 7 | 5.26e-07 |
| FC edges vs PANSS (partial corr) | IKEM | Surface IndiPar | PANSS panss_general_total (21 Yeo7 edges) | 21 | Partial corr (age/sex) using perm p (partial corr) + BH-FDR within dataset×pipeline×PANSS across 21 edges | q<=0.05 | 0 | 0.1054 |
| FC edges vs PANSS (partial corr) | IKEM | Surface IndiPar | PANSS panss_negative_total (21 Yeo7 edges) | 21 | Partial corr (age/sex) using perm p (partial corr) + BH-FDR within dataset×pipeline×PANSS across 21 edges | q<=0.05 | 0 | 0.2583 |
| FC edges vs PANSS (partial corr) | IKEM | Surface IndiPar | PANSS panss_positive_total (21 Yeo7 edges) | 21 | Partial corr (age/sex) using perm p (partial corr) + BH-FDR within dataset×pipeline×PANSS across 21 edges | q<=0.05 | 1 | 0.0126 |
| FC edges vs PANSS (partial corr) | IKEM | Surface IndiPar | PANSS panss_total (21 Yeo7 edges) | 21 | Partial corr (age/sex) using perm p (partial corr) + BH-FDR within dataset×pipeline×PANSS across 21 edges | q<=0.05 | 0 | 0.1092 |
| QPP event-rate AUC (Patient vs Control) | IKEM | Surface IndiPar | AUC of event rate (per subject) | 1 | Wilcoxon rank-sum (P vs C); no multiplicity (single test per dataset×pipeline) | p<=0.05 | 0 | 0.1468 |
| QPP event-rate AUC vs PANSS (partial corr) | IKEM | Surface IndiPar | PANSS panss_general_total | 1 | Partial corr (age/sex) using perm p (partial corr); no multiplicity (single test per dataset×pipeline×PANSS) | p<=0.05 | 1 | 0.018 |
| QPP event-rate AUC vs PANSS (partial corr) | IKEM | Surface IndiPar | PANSS panss_negative_total | 1 | Partial corr (age/sex) using perm p (partial corr); no multiplicity (single test per dataset×pipeline×PANSS) | p<=0.05 | 1 | 0.0438 |
| QPP event-rate AUC vs PANSS (partial corr) | IKEM | Surface IndiPar | PANSS panss_positive_total | 1 | Partial corr (age/sex) using perm p (partial corr); no multiplicity (single test per dataset×pipeline×PANSS) | p<=0.05 | 0 | 0.1356 |
| DMN–DAN opposition (dmndan_opp) | IKEM_vs_NUDZ | Surface IndiPar | DMN–DAN opposition (dmndan_opp) | 1 | Wilcoxon rank-sum comparing IKEM vs NUDZ within pipeline; no multiplicity | p<=0.05 | 1 | 0.047 |
| QPP rank distribution | IKEM_vs_NUDZ | Surface IndiPar | Selected component rank distribution (1/2/3) | 1 | Chi-square homogeneity (2x3) comparing IKEM vs NUDZ within pipeline; no multiplicity | p<=0.05 | 1 | 0.00727 |
| Delta FC (Patient - Control) | NUDZ | Surface IndiPar | Yeo7 between-network FC (z); 21 edges | 21 | Welch t-test (edgewise) + BH-FDR within dataset×pipeline across 21 edges | q<=0.05 | 9 | 4.092e-05 |
| FC edges vs PANSS (partial corr) | NUDZ | Surface IndiPar | PANSS panss_general_total (21 Yeo7 edges) | 21 | Partial corr (age/sex) using perm p (partial corr) + BH-FDR within dataset×pipeline×PANSS across 21 edges | q<=0.05 | 0 | 0.6488 |
| FC edges vs PANSS (partial corr) | NUDZ | Surface IndiPar | PANSS panss_negative_total (21 Yeo7 edges) | 21 | Partial corr (age/sex) using perm p (partial corr) + BH-FDR within dataset×pipeline×PANSS across 21 edges | q<=0.05 | 0 | 0.3801 |
| FC edges vs PANSS (partial corr) | NUDZ | Surface IndiPar | PANSS panss_positive_total (21 Yeo7 edges) | 21 | Partial corr (age/sex) using perm p (partial corr) + BH-FDR within dataset×pipeline×PANSS across 21 edges | q<=0.05 | 0 | 0.2562 |
| FC edges vs PANSS (partial corr) | NUDZ | Surface IndiPar | PANSS panss_total (21 Yeo7 edges) | 21 | Partial corr (age/sex) using perm p (partial corr) + BH-FDR within dataset×pipeline×PANSS across 21 edges | q<=0.05 | 0 | 0.4265 |
| QPP event-rate AUC (Patient vs Control) | NUDZ | Surface IndiPar | AUC of event rate (per subject) | 1 | Wilcoxon rank-sum (P vs C); no multiplicity (single test per dataset×pipeline) | p<=0.05 | 1 | 0.01838 |
| QPP event-rate AUC vs PANSS (partial corr) | NUDZ | Surface IndiPar | PANSS panss_general_total | 1 | Partial corr (age/sex) using perm p (partial corr); no multiplicity (single test per dataset×pipeline×PANSS) | p<=0.05 | 0 | 0.631 |
| QPP event-rate AUC vs PANSS (partial corr) | NUDZ | Surface IndiPar | PANSS panss_negative_total | 1 | Partial corr (age/sex) using perm p (partial corr); no multiplicity (single test per dataset×pipeline×PANSS) | p<=0.05 | 0 | 0.9651 |
| QPP event-rate AUC vs PANSS (partial corr) | NUDZ | Surface IndiPar | PANSS panss_positive_total | 1 | Partial corr (age/sex) using perm p (partial corr); no multiplicity (single test per dataset×pipeline×PANSS) | p<=0.05 | 0 | 0.7215 |

### Supplementary Table S1 (Page 2 of 4)

| analysis | dataset | pipeline | measure | family_size | test_and_correction | threshold | n_significant | min_p_or_q |
|---|---|---|---|---|---|---|---|---|
| QPP rank distribution | IKEM_vs_NUDZ | Surface Schaefer200 | Selected component rank distribution (1/2/3) | 1 | Chi-square homogeneity (2x3) comparing IKEM vs NUDZ within pipeline; no multiplicity | p<=0.05 | 0 | 0.07817 |
| Delta FC (Patient - Control) | NUDZ | Surface Schaefer200 | Yeo7 between-network FC (z); 21 edges | 21 | Welch t-test (edgewise) + BH-FDR within dataset×pipeline across 21 edges | q<=0.05 | 8 | 5.232e-05 |
| FC edges vs PANSS (partial corr) | NUDZ | Surface Schaefer200 | PANSS panss_general_total (21 Yeo7 edges) | 21 | Partial corr (age/sex) using perm p (partial corr) + BH-FDR within dataset×pipeline×PANSS across 21 edges | q<=0.05 | 0 | 0.7037 |
| FC edges vs PANSS (partial corr) | NUDZ | Surface Schaefer200 | PANSS panss_negative_total (21 Yeo7 edges) | 21 | Partial corr (age/sex) using perm p (partial corr) + BH-FDR within dataset×pipeline×PANSS across 21 edges | q<=0.05 | 0 | 0.315 |
| FC edges vs PANSS (partial corr) | NUDZ | Surface Schaefer200 | PANSS panss_positive_total (21 Yeo7 edges) | 21 | Partial corr (age/sex) using perm p (partial corr) + BH-FDR within dataset×pipeline×PANSS across 21 edges | q<=0.05 | 0 | 0.8173 |
| FC edges vs PANSS (partial corr) | NUDZ | Surface Schaefer200 | PANSS panss_total (21 Yeo7 edges) | 21 | Partial corr (age/sex) using perm p (partial corr) + BH-FDR within dataset×pipeline×PANSS across 21 edges | q<=0.05 | 0 | 0.4046 |
| QPP event-rate AUC (Patient vs Control) | NUDZ | Surface Schaefer200 | AUC of event rate (per subject) | 1 | Wilcoxon rank-sum (P vs C); no multiplicity (single test per dataset×pipeline) | p<=0.05 | 1 | 0.03218 |
| QPP event-rate AUC vs PANSS (partial corr) | NUDZ | Surface Schaefer200 | PANSS panss_general_total | 1 | Partial corr (age/sex) using perm p (partial corr); no multiplicity (single test per dataset×pipeline×PANSS) | p<=0.05 | 0 | 0.2099 |
| QPP event-rate AUC vs PANSS (partial corr) | NUDZ | Surface Schaefer200 | PANSS panss_negative_total | 1 | Partial corr (age/sex) using perm p (partial corr); no multiplicity (single test per dataset×pipeline×PANSS) | p<=0.05 | 0 | 0.156 |
| QPP event-rate AUC vs PANSS (partial corr) | NUDZ | Surface Schaefer200 | PANSS panss_positive_total | 1 | Partial corr (age/sex) using perm p (partial corr); no multiplicity (single test per dataset×pipeline×PANSS) | p<=0.05 | 0 | 0.5273 |
| Delta FC (Patient - Control) | IKEM | Vol Brainnetome246 | Yeo7 between-network FC (z); 21 edges | 21 | Welch t-test (edgewise) + BH-FDR within dataset×pipeline across 21 edges | q<=0.05 | 3 | 0.02719 |
| FC edges vs PANSS (partial corr) | IKEM | Vol Brainnetome246 | PANSS panss_general_total (21 Yeo7 edges) | 21 | Partial corr (age/sex) using perm p (partial corr) + BH-FDR within dataset×pipeline×PANSS across 21 edges | q<=0.05 | 0 | 0.2877 |
| FC edges vs PANSS (partial corr) | IKEM | Vol Brainnetome246 | PANSS panss_negative_total (21 Yeo7 edges) | 21 | Partial corr (age/sex) using perm p (partial corr) + BH-FDR within dataset×pipeline×PANSS across 21 edges | q<=0.05 | 0 | 0.1687 |
| FC edges vs PANSS (partial corr) | IKEM | Vol Brainnetome246 | PANSS panss_positive_total (21 Yeo7 edges) | 21 | Partial corr (age/sex) using perm p (partial corr) + BH-FDR within dataset×pipeline×PANSS across 21 edges | q<=0.05 | 0 | 0.8673 |

### Supplementary Table S1 (Page 3 of 4)

| analysis | dataset | pipeline | measure | family_size | test_and_correction | threshold | n_significant | min_p_or_q |
|---|---|---|---|---|---|---|---|---|
| Delta FC (Patient - Control) | IKEM | Vol Schaefer200 | Yeo7 between-network FC (z); 21 edges | 21 | Welch t-test (edgewise) + BH-FDR within dataset×pipeline across 21 edges | q<=0.05 | 4 | 0.005546 |
| FC edges vs PANSS (partial corr) | IKEM | Vol Schaefer200 | PANSS panss_general_total (21 Yeo7 edges) | 21 | Partial corr (age/sex) using perm p (partial corr) + BH-FDR within dataset×pipeline×PANSS across 21 edges | q<=0.05 | 0 | 0.4995 |
| FC edges vs PANSS (partial corr) | IKEM | Vol Schaefer200 | PANSS panss_negative_total (21 Yeo7 edges) | 21 | Partial corr (age/sex) using perm p (partial corr) + BH-FDR within dataset×pipeline×PANSS across 21 edges | q<=0.05 | 0 | 0.2305 |
| FC edges vs PANSS (partial corr) | IKEM | Vol Schaefer200 | PANSS panss_positive_total (21 Yeo7 edges) | 21 | Partial corr (age/sex) using perm p (partial corr) + BH-FDR within dataset×pipeline×PANSS across 21 edges | q<=0.05 | 0 | 0.8697 |
| FC edges vs PANSS (partial corr) | IKEM | Vol Schaefer200 | PANSS panss_total (21 Yeo7 edges) | 21 | Partial corr (age/sex) using perm p (partial corr) + BH-FDR within dataset×pipeline×PANSS across 21 edges | q<=0.05 | 0 | 0.3549 |
| QPP event-rate AUC (Patient vs Control) | IKEM | Vol Schaefer200 | AUC of event rate (per subject) | 1 | Wilcoxon rank-sum (P vs C); no multiplicity (single test per dataset×pipeline) | p<=0.05 | 0 | 0.9958 |
| QPP event-rate AUC vs PANSS (partial corr) | IKEM | Vol Schaefer200 | PANSS panss_general_total | 1 | Partial corr (age/sex) using perm p (partial corr); no multiplicity (single test per dataset×pipeline×PANSS) | p<=0.05 | 0 | 0.201 |
| QPP event-rate AUC vs PANSS (partial corr) | IKEM | Vol Schaefer200 | PANSS panss_negative_total | 1 | Partial corr (age/sex) using perm p (partial corr); no multiplicity (single test per dataset×pipeline×PANSS) | p<=0.05 | 1 | 0.041 |
| QPP event-rate AUC vs PANSS (partial corr) | IKEM | Vol Schaefer200 | PANSS panss_positive_total | 1 | Partial corr (age/sex) using perm p (partial corr); no multiplicity (single test per dataset×pipeline×PANSS) | p<=0.05 | 0 | 0.05499 |
| DMN–DAN opposition (dmndan_opp) | IKEM_vs_NUDZ | Vol Schaefer200 | DMN–DAN opposition (dmndan_opp) | 1 | Wilcoxon rank-sum comparing IKEM vs NUDZ within pipeline; no multiplicity | p<=0.05 | 0 | 0.07518 |
| QPP rank distribution | IKEM_vs_NUDZ | Vol Schaefer200 | Selected component rank distribution (1/2/3) | 1 | Chi-square homogeneity (2x3) comparing IKEM vs NUDZ within pipeline; no multiplicity | p<=0.05 | 0 | 0.4185 |
| Delta FC (Patient - Control) | NUDZ | Vol Schaefer200 | Yeo7 between-network FC (z); 21 edges | 21 | Welch t-test (edgewise) + BH-FDR within dataset×pipeline across 21 edges | q<=0.05 | 6 | 0.002432 |
| FC edges vs PANSS (partial corr) | NUDZ | Vol Schaefer200 | PANSS panss_general_total (21 Yeo7 edges) | 21 | Partial corr (age/sex) using perm p (partial corr) + BH-FDR within dataset×pipeline×PANSS across 21 edges | q<=0.05 | 0 | 0.6178 |
| FC edges vs PANSS (partial corr) | NUDZ | Vol Schaefer200 | PANSS panss_negative_total (21 Yeo7 edges) | 21 | Partial corr (age/sex) using perm p (partial corr) + BH-FDR within dataset×pipeline×PANSS across 21 edges | q<=0.05 | 0 | 0.4449 |
| FC edges vs PANSS (partial corr) | NUDZ | Vol Schaefer200 | PANSS panss_positive_total (21 Yeo7 edges) | 21 | Partial corr (age/sex) using perm p (partial corr) + BH-FDR within dataset×pipeline×PANSS across 21 edges | q<=0.05 | 0 | 0.5543 |
| FC edges vs PANSS (partial corr) | NUDZ | Vol Schaefer200 | PANSS panss_total (21 Yeo7 edges) | 21 | Partial corr (age/sex) using perm p (partial corr) + BH-FDR within dataset×pipeline×PANSS across 21 edges | q<=0.05 | 0 | 0.4628 |
| QPP event-rate AUC (Patient vs Control) | NUDZ | Vol Schaefer200 | AUC of event rate (per subject) | 1 | Wilcoxon rank-sum (P vs C); no multiplicity (single test per dataset×pipeline) | p<=0.05 | 0 | 0.4779 |

### Supplementary Table S1 (Page 4 of 4)

| analysis | dataset | pipeline | measure | family_size | test_and_correction | threshold | n_significant | min_p_or_q |
|---|---|---|---|---|---|---|---|---|
| QPP rank distribution | IKEM | Surface IndiPar vs Vol Brainnetome246 | Selected component rank distribution (1/2/3) | 1 | Chi-square homogeneity (2x3); no multiplicity (single pairwise comparison) | p<=0.05 | 1 | 2.22e-16 |
| QPP rank distribution | IKEM | Surface IndiPar vs Vol Schaefer200 | Selected component rank distribution (1/2/3) | 1 | Chi-square homogeneity (2x3); no multiplicity (single pairwise comparison) | p<=0.05 | 1 | 0 |
| QPP rank distribution | IKEM | Surface Schaefer200 vs Vol Brainnetome246 | Selected component rank distribution (1/2/3) | 1 | Chi-square homogeneity (2x3); no multiplicity (single pairwise comparison) | p<=0.05 | 1 | 1.636e-07 |
| QPP rank distribution | IKEM | Surface Schaefer200 vs Vol Schaefer200 | Selected component rank distribution (1/2/3) | 1 | Chi-square homogeneity (2x3); no multiplicity (single pairwise comparison) | p<=0.05 | 1 | 2.614e-09 |
| QPP rank distribution | IKEM | Vol Brainnetome246 vs Vol Schaefer200 | Selected component rank distribution (1/2/3) | 1 | Chi-square homogeneity (2x3); no multiplicity (single pairwise comparison) | p<=0.05 | 0 | 0.1886 |
| DMN–DAN opposition (dmndan_opp) | NUDZ | Surface IndiPar vs Surface Schaefer200 | DMN–DAN opposition (dmndan_opp) | 1 | Wilcoxon rank-sum; no multiplicity (single pairwise comparison) | p<=0.05 | 1 | 3.057e-41 |
| DMN–DAN opposition (dmndan_opp) | NUDZ | Surface IndiPar vs Vol Brainnetome246 | DMN–DAN opposition (dmndan_opp) | 1 | Wilcoxon rank-sum; no multiplicity (single pairwise comparison) | p<=0.05 | 1 | 1.154e-65 |
| DMN–DAN opposition (dmndan_opp) | NUDZ | Surface IndiPar vs Vol Schaefer200 | DMN–DAN opposition (dmndan_opp) | 1 | Wilcoxon rank-sum; no multiplicity (single pairwise comparison) | p<=0.05 | 1 | 1.687e-43 |
| DMN–DAN opposition (dmndan_opp) | NUDZ | Surface Schaefer200 vs Vol Brainnetome246 | DMN–DAN opposition (dmndan_opp) | 1 | Wilcoxon rank-sum; no multiplicity (single pairwise comparison) | p<=0.05 | 1 | 9.115e-20 |
| DMN–DAN opposition (dmndan_opp) | NUDZ | Surface Schaefer200 vs Vol Schaefer200 | DMN–DAN opposition (dmndan_opp) | 1 | Wilcoxon rank-sum; no multiplicity (single pairwise comparison) | p<=0.05 | 0 | 0.6979 |
| DMN–DAN opposition (dmndan_opp) | NUDZ | Vol Brainnetome246 vs Vol Schaefer200 | DMN–DAN opposition (dmndan_opp) | 1 | Wilcoxon rank-sum; no multiplicity (single pairwise comparison) | p<=0.05 | 1 | 1.74e-19 |
| QPP rank distribution | NUDZ | Surface IndiPar vs Surface Schaefer200 | Rank-1 prevalence (rank==1) | 1 | Chi-square (2x2: rank1 vs non-rank1); no multiplicity (single pairwise comparison) | p<=0.05 | 1 | 0.02315 |
| QPP rank distribution | NUDZ | Surface IndiPar vs Vol Brainnetome246 | Rank-1 prevalence (rank==1) | 1 | Chi-square (2x2: rank1 vs non-rank1); no multiplicity (single pairwise comparison) | p<=0.05 | 1 | 0 |
| QPP rank distribution | NUDZ | Surface IndiPar vs Vol Schaefer200 | Rank-1 prevalence (rank==1) | 1 | Chi-square (2x2: rank1 vs non-rank1); no multiplicity (single pairwise comparison) | p<=0.05 | 1 | 0 |
| QPP rank distribution | NUDZ | Surface IndiPar vs Surface Schaefer200 | Selected component rank distribution (1/2/3) | 1 | Chi-square homogeneity (2x3); no multiplicity (single pairwise comparison) | p<=0.05 | 1 | 0.004353 |
| QPP rank distribution | NUDZ | Surface IndiPar vs Vol Brainnetome246 | Selected component rank distribution (1/2/3) | 1 | Chi-square homogeneity (2x3); no multiplicity (single pairwise comparison) | p<=0.05 | 1 | 0 |
| QPP rank distribution | NUDZ | Surface IndiPar vs Vol Schaefer200 | Selected component rank distribution (1/2/3) | 1 | Chi-square homogeneity (2x3); no multiplicity (single pairwise comparison) | p<=0.05 | 1 | 0 |
| QPP rank distribution | NUDZ | Surface Schaefer200 vs Vol Brainnetome246 | Selected component rank distribution (1/2/3) | 1 | Chi-square homogeneity (2x3); no multiplicity (single pairwise comparison) | p<=0.05 | 1 | 4.408e-14 |
| QPP rank distribution | NUDZ | Surface Schaefer200 vs Vol Schaefer200 | Selected component rank distribution (1/2/3) | 1 | Chi-square homogeneity (2x3); no multiplicity (single pairwise comparison) | p<=0.05 | 1 | 3.36e-13 |
| QPP rank distribution | NUDZ | Vol Brainnetome246 vs Vol Schaefer200 | Selected component rank distribution (1/2/3) | 1 | Chi-square homogeneity (2x3); no multiplicity (single pairwise comparison) | p<=0.05 | 0 | 0.3433 |
